# Loss of Complement Factor H impairs antioxidant capacity and energy metabolism of human RPE cells

**DOI:** 10.1101/2020.01.08.898551

**Authors:** Angela Armento, Sabina Honisch, Vasiliki Panagiotakopoulou, Inga Sonntag, Anke Jacob, Ellen Kilger, Michela Deleidi, Simon Clark, Marius Ueffing

**Affiliations:** Institute for Ophthalmic Research, Department for Ophthalmology, Tübingen, Germany; German Center for Neurodegenerative Diseases (DZNE), Tübingen, Germany; Hertie-Institute for Clinical Brain Research, University of Tübingen, Tübingen, Germany

**Keywords:** Age-related macular degeneration (AMD), complement factor H (CFH), retinal pigment epithelium (RPE), glycolysis, mitochondria respiration, oxidative stress

## Abstract

Age-related macular degeneration (AMD) is the leading cause of blindness in the elderly population. About 50% of AMD patients present polymorphisms in the Complement Factor H (*CFH*) gene, coding for Factor H protein (FH). AMD-associated *CFH* risk variants, Y402H in particular, impair FH function leading to complement overactivation. In AMD, retinal homeostasis is compromised due to dysfunction of retinal pigment epithelium (RPE) cells. Whether FH contributes to AMD pathogenesis only via complement system dysregulation remains unclear. To investigate the potential role of FH on energy metabolism and oxidative stress in RPE cells, we silenced *CFH* in human hTERT-RPE1 cells. FH-deprived RPE cells exposed to oxidative insult, showed altered metabolic homeostasis, including reduction of glycolysis and mitochondrial respiration, paralleled by an increase in lipid peroxidation. Our data suggest that FH protects RPE cells from oxidative stress and metabolic reprogramming, highlighting a novel function for FH in AMD pathogenesis.

**Graphical abstract:** 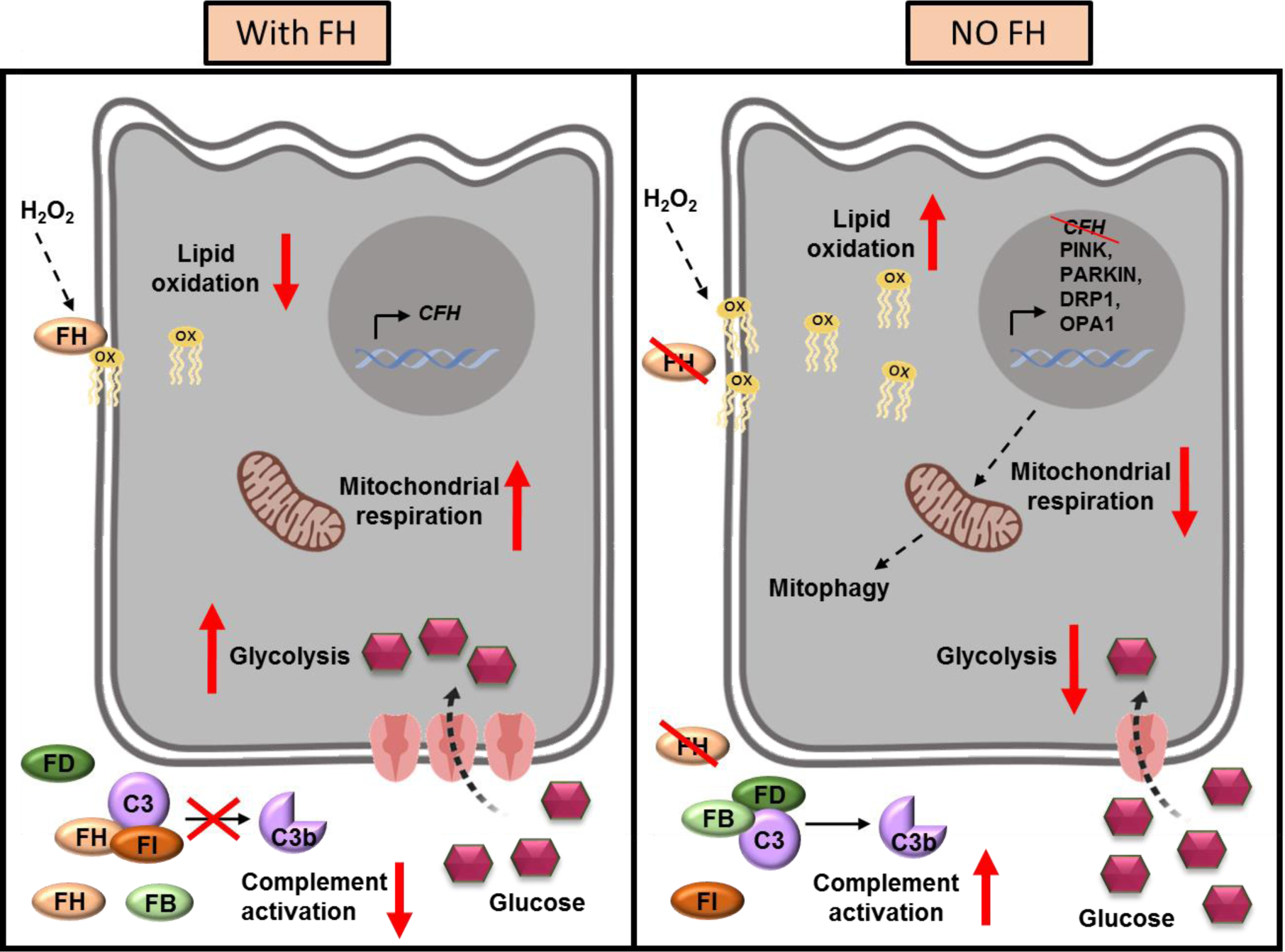

## Introduction

Age-related macular degeneration (AMD) is characterized by a progressive degeneration of the macula, leading to central vision loss and ultimately blindness. AMD is a complex disease, involving ageing, genetic predisposition and environmental factors, and a full understanding of AMD pathogenesis is lacking, which makes drug discovery challenging. AMD affects mainly the elderly population and it is estimated that around 200 million people will be affected by 2020 and 300 million by 2040 [1]. The clinical classification of AMD is based on the appearance of the retina in fundus imaging, and two forms of AMD are defined: “wet” and “dry” AMD. Fundus photographs showing traits of neovascularization are distinctive of “wet” AMD. Indeed, wet AMD is characterized by the formation of new and unorganized blood vessels network which invade the Bruch’s membrane (BM) and RPE layer, damaging the retina, a process termed choroidal neovascularisation (CNV). Wet AMD displays a more severe phenotype, affects circa 10-15% of AMD patients and anti-VEGF therapy can be applied to slow down the progression of the disease. Dry AMD, for which no therapy is applicable, affects the majority of AMD patients and it is clinically defined by the appearance of areas of geographic atrophy with borders of hyperpigmented RPE cells, as seen in fundus images [2]. Dry AMD is defined by damage and degeneration of RPE cells, leading to photoreceptors malfunctioning and visual loss. Early stages of both AMD forms are characterized by the presence of deposits, called drusen, between the BM and the RPE layer [3]. An intact and well-functioning RPE cell layer, which provides a barrier between the neuroretina and the choroid capillary network, is essential for the maintenance of retinal homeostasis. In the presence of drusen or altered extracellular matrix (ECM) of BM, functionality of RPE cells may be impaired [4]. In addition, RPE cells fulfil several key functions, such as phagocytosis of the photoreceptor outer segments, transport of nutrients, preservation of the retinal structure and, most importantly, due to their high antioxidant capacity, RPE cells protect the retina from photo-oxidation and oxidative damage [5]. Recently, evidence in support of the hypothesis that a bioenergetic failure of RPE cells may be at the basis for AMD pathology has been provided [6]. The retinal microenvironment is highly oxidized due to a very high energy demand and photo-oxidation. Ageing processes in combination with external stressors, such as smoking or a high fat diet [7, 8], force RPE cells to deal with excessive levels of oxidative stress. In fact, energy metabolism of primary RPE cells isolated from AMD patients was found to be strongly impaired compared to RPE cells from healthy controls [9]. In this model, glycolysis as well as mitochondrial respiration were reduced in RPE cells from AMD patients [9]. Other studies showed that mitochondrial dysfunction in RPE cells may represent a relevant AMD feature. Of the proteins differentially expressed in RPE cells from AMD donors and healthy controls, many are mitochondrial proteins [10]. Moreover, it has been shown in a mouse model that conditionally-induced mitochondrial damage in RPE cells leads to cell metabolic reprogramming and photoreceptors malfunction [11].

So far, there is lack of knowledge on the impact of the high-risk genetic variants on RPE cells homeostasis. A large portion of AMD genetic high-risk variants is located in genes coding for complement system regulatory proteins (FH, FI, C3, FB/C2) [12]. In particular, a common polymorphism in the complement factor H (*CFH*) gene, leads to an amino acid exchange from a tyrosine to histidine at position 402 (Y402H) in the two proteins encoded by the gene: factor H (FH) and its truncated form factor H-like protein 1 (FHL-1) [13]. This common polymorphism is strongly associated with increased risk for AMD and may account for ∼50% of AMD cases in the United States [14]. The mechanism by which the FH H402 variant confers predisposition for AMD is not clear. The FH H402 variant impairs FH and FHL-1 function, leading to uncontrolled complement system activation *in vitro* [15]. Recent studies highlighted the possibility that FH is not only contributing to AMD through its classical role in complement regulation, but it may influence other processes. For example, it has been shown that the FH H402 variant presents altered binding affinity to C-reactive protein or malondialdehyde, indicating a possible role in inflammation and lipoprotein degradation [16, 17], processes both associated with AMD pathogenesis [18, 19]. These defects may influence the ability of RPE cells to cope with oxidative stress. This study was designed to unravel the impact of endogenous FH loss on RPE cells metabolism and their vulnerability toward oxidative stress. We employed RNA interference to decrease FH levels in the hTERT-RPE1 human cell line. Perhaps unsurprisingly, FH reduction led to activation of the complement system. Using the Seahorse Extracellular Flux Analyzer to measure bioenergetics, we observed that knock-down of the *CFH* gene negatively affects mitochondrial and glycolytic function of RPE cells when compared to controls. The impairment was even more pronounced when cells were exposed to oxidative stress by pre-treatment with hydrogen peroxide. The effects of FH reduction on energy metabolism were accompanied by transcriptional regulation of several glucose metabolism genes as well as genes modulating mitochondrial stability. RPE cells lacking FH showed a significant increase in lipid peroxidation, which is a key aspect of AMD pathogenesis and, in parallel, cell viability was decreased. Our results suggest that endogenous FH, produced by RPE cells, not only modulates the extracellular microenvironment via its negative effect on complement activation, but also has an intracellular impact on the antioxidant functions and metabolic homeostasis of RPE cells, refining the knowledge on how FH is involved in AMD processes.

## RESULTS

### FH reduction leads to complement activation in RPE cells

To investigate the role of FH, we used siRNA to silence the *CFH* gene in hTERT-RPE1 established cell lines and we induced a mild oxidative stress through hydrogen peroxide pre-treatment (200 µM for 90 minutes). This set-up provides the chance to study *in vitro* the combination of endogenous FH dysregulation and environmental factors contributing to AMD, which increase oxidative stress. We monitored the efficiency of *CFH* silencing in all experimental conditions, including PBS and H_2_O_2_ pre-treated cells after 48 hours in culture. Significantly reduced *CFH* mRNA was detected in *CFH* knock-down cells compared to the siNeg control cells, achieving almost 90% silencing of the *CFH* gene (Fig 1A). The FH protein was almost undetected in cell culture supernatants collected at the same time point from the si*CFH* cells compared to controls (Fig 1B). Based on gene expression levels of RPE markers: Bestrophin 1 (BEST1), Retinoid Isomerohydrolase (RPE65) and Tight junction protein ZO-1 (TJP1), RPE characteristics in experimental conditions were not altered (Supplementary Fig 1). Depletion of the FH protein led to reduced regulation of complement activation. C3b is the first cleaved peptide triggering complement activation and denotes the amplification loop of the proteolytic cascade, characteristic of the alternative pathway of the complement system [20]. Therefore, we assessed secreted levels of C3b in the *CFH* knock-down cells by both ELISA and Western blot and found in both cases a 2-fold increase in detectable C3b (Fig. 1C-D).

**Figure 1.**
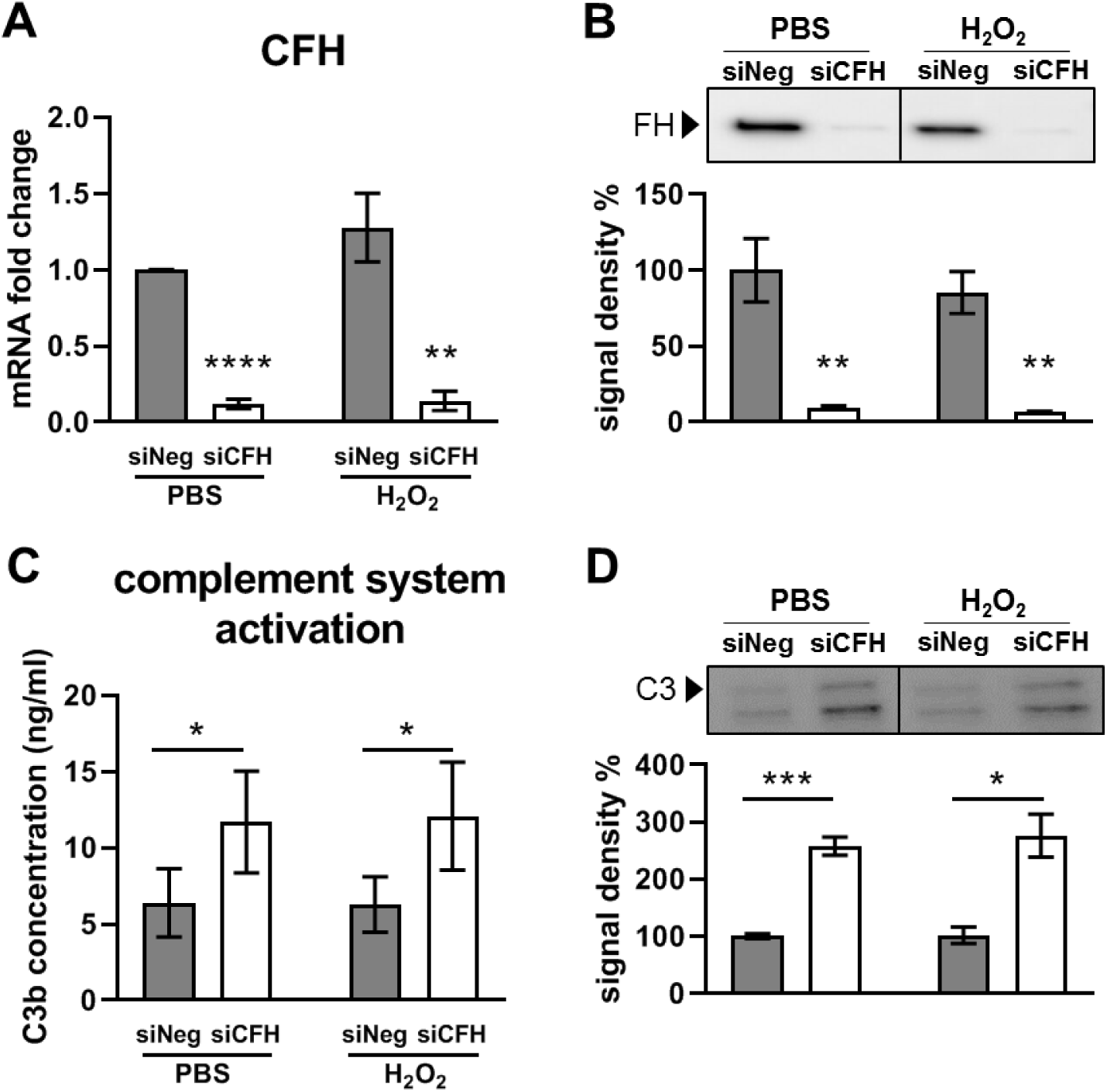
FH reduction leads to complement system activation in RPE cells. hTERT-RPE1 cells were seeded, let attach overnight and silenced for 24 hours with negative control (siNeg) or *CFH* specific (si*CFH*) siRNA. Cells were exposed for 90 minutes to 200 µM H_2_O_2_ or PBS and cell pellets and cell culture supernatants were collected for further processing after 48 hours. **A** Monitoring of *CFH* expression by qRT-PCR analyses in silencing negative control (siNeg) and specific *CFH* silenced (si*CFH*) hTERT-RPE1 cells. Data are normalized to housekeeping gene PRPL0 using Δ ΔCt method. SEM is shown, n=3. **B** Western blot analyses of FH protein levels in cell culture supernatants of hTERT-RPE1 in the same conditions as A. Quantification of signal density of 4 independent experiments is shown. **C** C3b ELISA analyses of cell culture supernatants of hTERT-RPE1 cells. SEM is shown, n=4. **D** Western blot analyses of C3 protein levels in cell culture supernatants of hTERT-RPE1 cells. Quantification of signal density of 3 independent experiments is shown. Significance was assessed by Student T-test. *p<0.05, **p<0.01, *** p<0.001, ****p<0.0001.

### FH loss increases vulnerability toward oxidative stress in RPE cells

In order to assess whether the silencing of *CFH* altered the response of hTERT-RPE1 cells to oxidative stress, we investigated cell lipid peroxidation levels after H_2_O_2_ treatment (Fig. 2A). In our model, lipid peroxidation levels were significantly increased only in the absence of FH 48 hours after the oxidative treatment (Fig. 2A). As shown in Fig 2B, cell viability was not affected in the absence of *CFH* expression in PBS alone, and pre-treatment with H_2_O_2_ had no effects on the siNeg control cells, confirming the known high antioxidant capacity of RPE cells [21]. However, cell viability was significantly reduced exclusively when RPE cells missing *CFH* expression were stimulated with H_2_O_2_ (Fig. 2B), indicating increased vulnerability toward a short exposure to oxidative stress in FH deprived RPE cells. In parallel, we investigated cell membrane damage via a cytotoxicity assay. Silencing of *CFH* in RPE cells led to an increase in RPE cell damage, irrespective of H_2_O_2_-induced oxidative stress (Fig. 2C).

**Figure 2.**
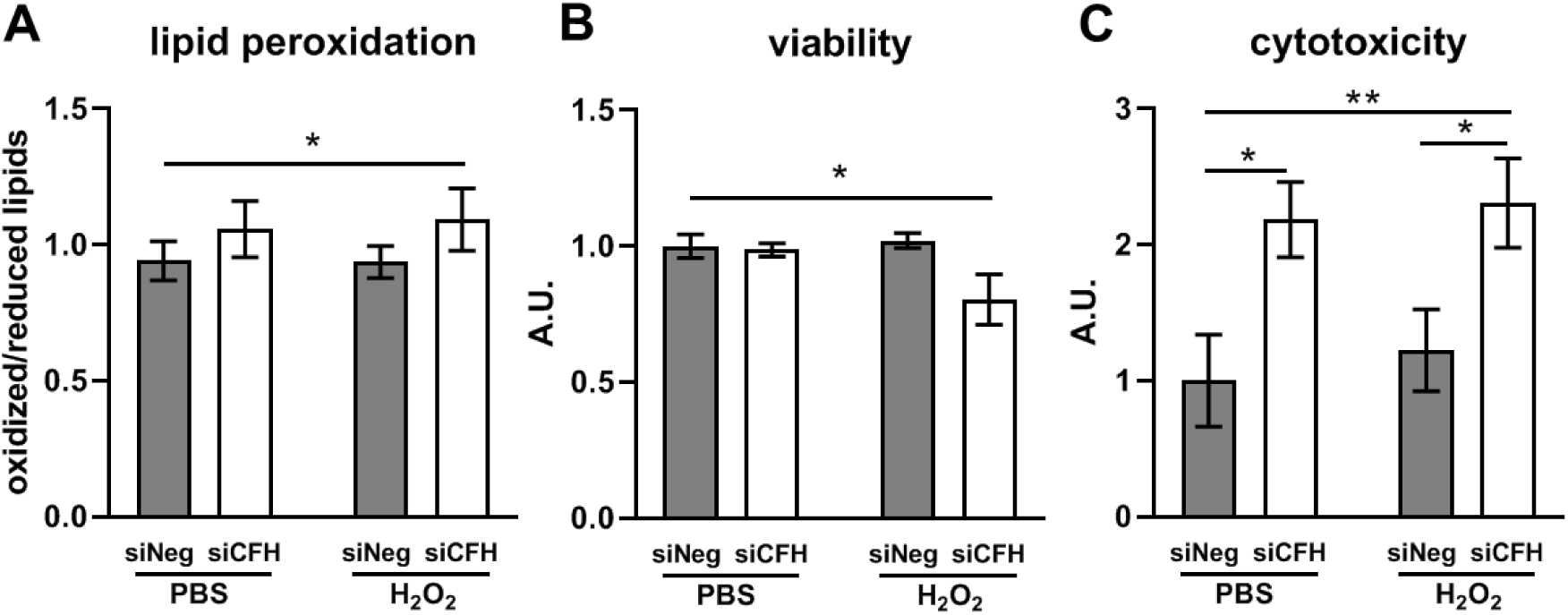
FH loss increases vulnerability toward oxidative stress RPE cells. hTERT-RPE1 cells were seeded, let attach overnight and silenced for 24 hours with negative control (siNeg) or *CFH* specific (si*CFH*) siRNA. Cells were exposed for 90 minutes to 200 µM H_2_O_2_ or PBS and specific dyes were added after 48 hours. **A** Lipid peroxidation levels were assessed via BODIPY® 581/591 C11 fluorescent dye. Fluorescence shift was measured at ∼590 nm and ∼510 nm. Data are shown as ratio oxidized/reduced lipids, higher bars indicate higher lipid peroxidation levels. SEM is shown, n=7 **B** Viability was assessed by cell-permeable fluorescent dye GF-AFC (glycyl-phenylalanyl-aminofluorocoumarin). SEM is shown, n=5 **C** Cytotoxicity levels were assessed by cell-impermeable fluorescent dye bis-AAF-R110. SEM is shown, n=5. A.U. arbitrary units. Significance was assessed by Student t-test (single effect) and two-way ANOVA (combined effects) as described in the methods section. * p<0.05, **p<0.01.

### FH loss impairs glycolysis in RPE cells

To investigate the influence of FH on RPE cell metabolism, the extracellular acidification rate (ECAR) was monitored as an indication of glycolytic function using the glycolysis stress test. Fig 3A shows a schematic representation of glycolysis pathway highlighting the substances used in the Seahorse analyses. Glucose, oligomycin and 2-deoxyglucose (2-DG) were sequentially injected (as shown by the arrows in Fig. 3B) to modulate glycolysis responses and ECAR. Fig. 3B shows ECAR measurements in siNeg cells and si*CFH* cells pretreated with PBS or 200 µM H_2_O_2_ for 90 minutes. Basal levels of glycolysis were found to be significantly lower by 43% in RPE cells deprived of FH, compared to siNeg controls (Fig 3C-D). This reduction was even more pronounced when the *CFH* knock-down cells were pre-treated with H_2_O_2_ (Fig 3C), with glycolysis being reduced by 63% compared to cells treated only with H_2_O_2_ (Fig 3C). Glycolytic capacity was significantly reduced only when *CFH* knock-down cells were pre-treated with H_2_O_2_ (Fig 3C), showing a reduction of 50 % compared to siNeg control cells (Fig 3D). Also, glycolytic reserve in si*CFH* H_2_O_2_ treated cells was slightly reduced compared to both siNeg controls (Fig 3E). Consistently, glucose uptake was reduced significantly in si*CFH* cells after H_2_O_2_ exposure compared to siNeg control cells (Fig 4A). In parallel, mRNA expression of glucose transporter GLUT1 was reduced in H_2_O_2_-treated si*CFH* cells compared to both treated and untreated controls (Fig 4B). Gene expression of LDHA (lactate dehydrogenase A), an isoform of LDH which preferentially converts pyruvate to lactate [22], was also significantly reduced in all si*CFH* cells, more pronouncedly after peroxide treatment (Fig 4B).

**Figure 3.**
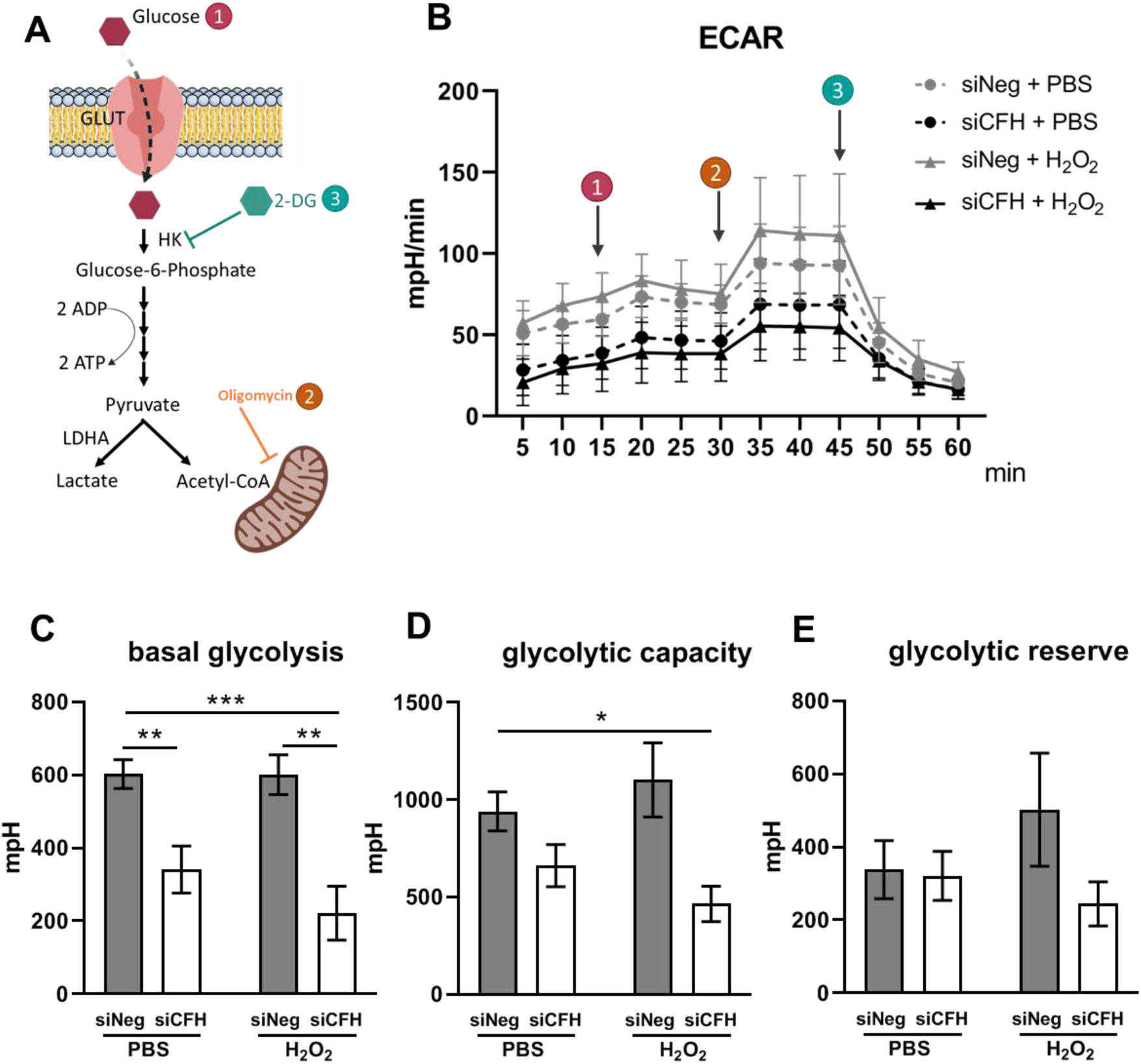
FH loss impairs glycolysis in RPE cells. **A** Schematic representation of glycolysis and steps targeted during Seahorse analyses (1,2,3) **B** hTERT-RPE1 cells were seeded, let attach overnight and silenced for 24 hours with negative control (siNeg) or *CFH* specific (si*CFH*) siRNA. 30.000 cells were transferred to Seahorse plates overnight and pre-treated for 90 minutes with 200 µM H_2_O_2_ or PBS. Curves show extracellular acidification rate (ECAR) measured after 48 hours. SEM is shown, n=4-8. Arrows indicate injection of glucose (1), oligomycin (2) and 2-deoxyglucose (2-DG,3). **C-E** Parameters of glycolytic function are calculated from data shown in B. Basal glycolysis (C), glycolytic capacity (D) and glycolytic reserve (E). Significance was assessed by Student t-test (single effect) and two-way ANOVA (combined effects) as described in the methods section. * p<0.05, **p<0.01, *** p<0.001

**Figure 4.**
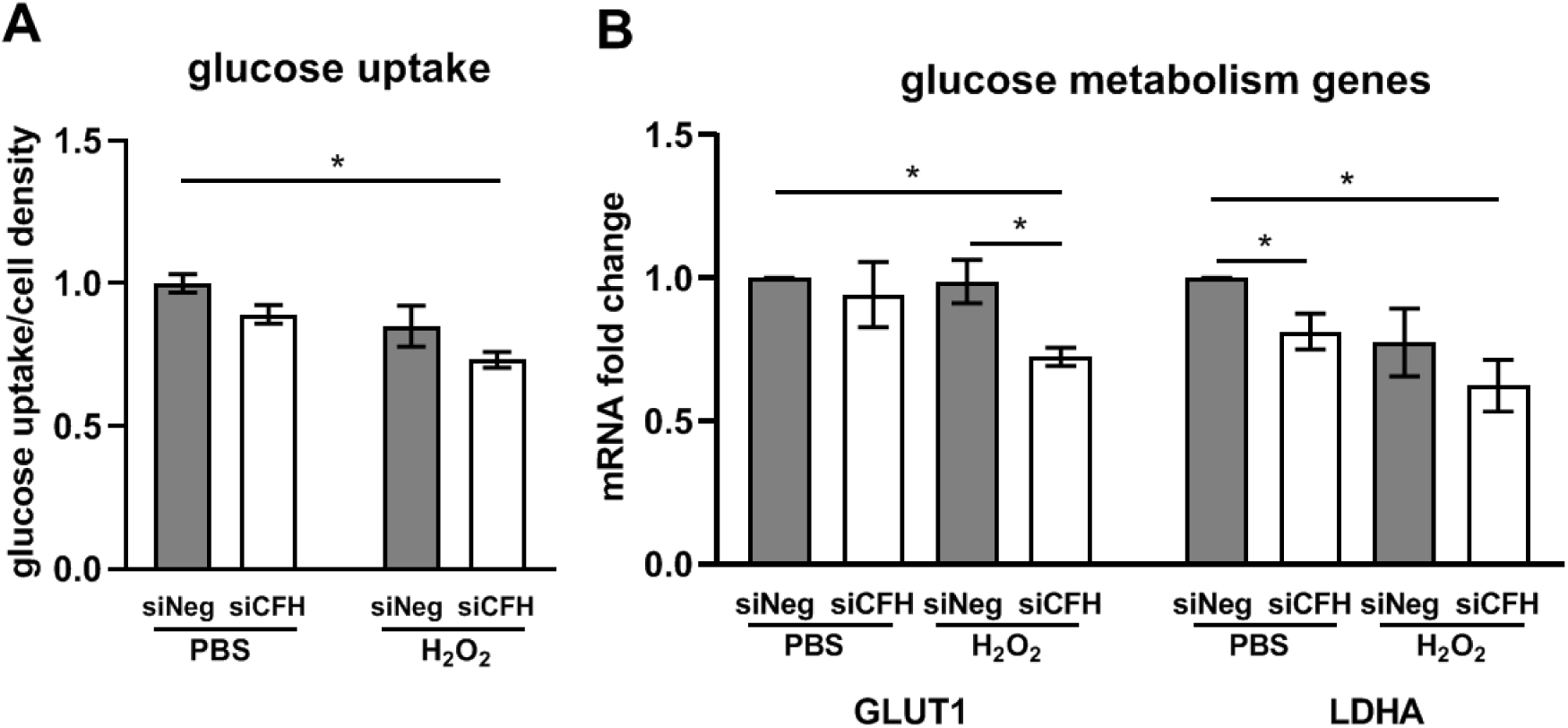
FH modulates glucose uptake and expression of glucose metabolism genes. hTERT-RPE1 cells were seeded, let attach overnight and silenced for 24 hours with negative control (siNeg) or *CFH* specific (si*CFH*) siRNA. Cells were exposed for 90 minutes to 200 µM H_2_O_2_ or PBS **A** Glucose uptake was measured 48 hours after H_2_O_2_ pre-treatment in siNeg control cells and in si*CFH* cells. SEM is shown, n=3. **B** gene expression analysis by qRT-PCR of glucose transporter 1 (GLUT1/SLC2A1) and glycolysis enzyme gene lactate dehydrogenase A (LDHA). SEM is shown, n=3. Data are normalized to housekeeping gene PRPL0 using Δ ΔCt method. Significance was assessed by Student t-test (single effect) and two-way ANOVA (combined effects) as described in the methods section. * p<0.05.

### FH loss impairs mitochondrial respiration in RPE cells

The potential influence of FH loss on mitochondrial respiration of RPE cells was assessed monitoring the oxygen consumption rate (OCR), an indication of mitochondria respiratory function using the cell mito stress test. Fig 5A shows a schematic representation of the oxidative phosphorylation (OxPhos) chain, highlighting the substances used in the Seahorse analyses. Oligomycin, carbonyl cyanide-4-(trifluoromethoxy)phenylhydrazone (FCCP) and antimycin/rotenone were sequentially injected (as shown by the arrows in Fig 5B) to assess OCR in different conditions and calculate parameters of mitochondrial function. Fig 5B shows OCR measurements in siNeg cells and si*CFH* cells pretreated with PBS or 200 µM H_2_O_2_ for 90 minutes. All parameters of mitochondrial respiration showed a clear trend of reduction in all si*CFH* groups (Fig 5C-D-E). However, the maximal respiration was significantly reduced in the absence of FH by 52 % in PBS-treated cells (Fig 5D). A slight increase in maximal respiration was observed when control cells were treated only with peroxide (Fig 5D).

**Figure 5.**
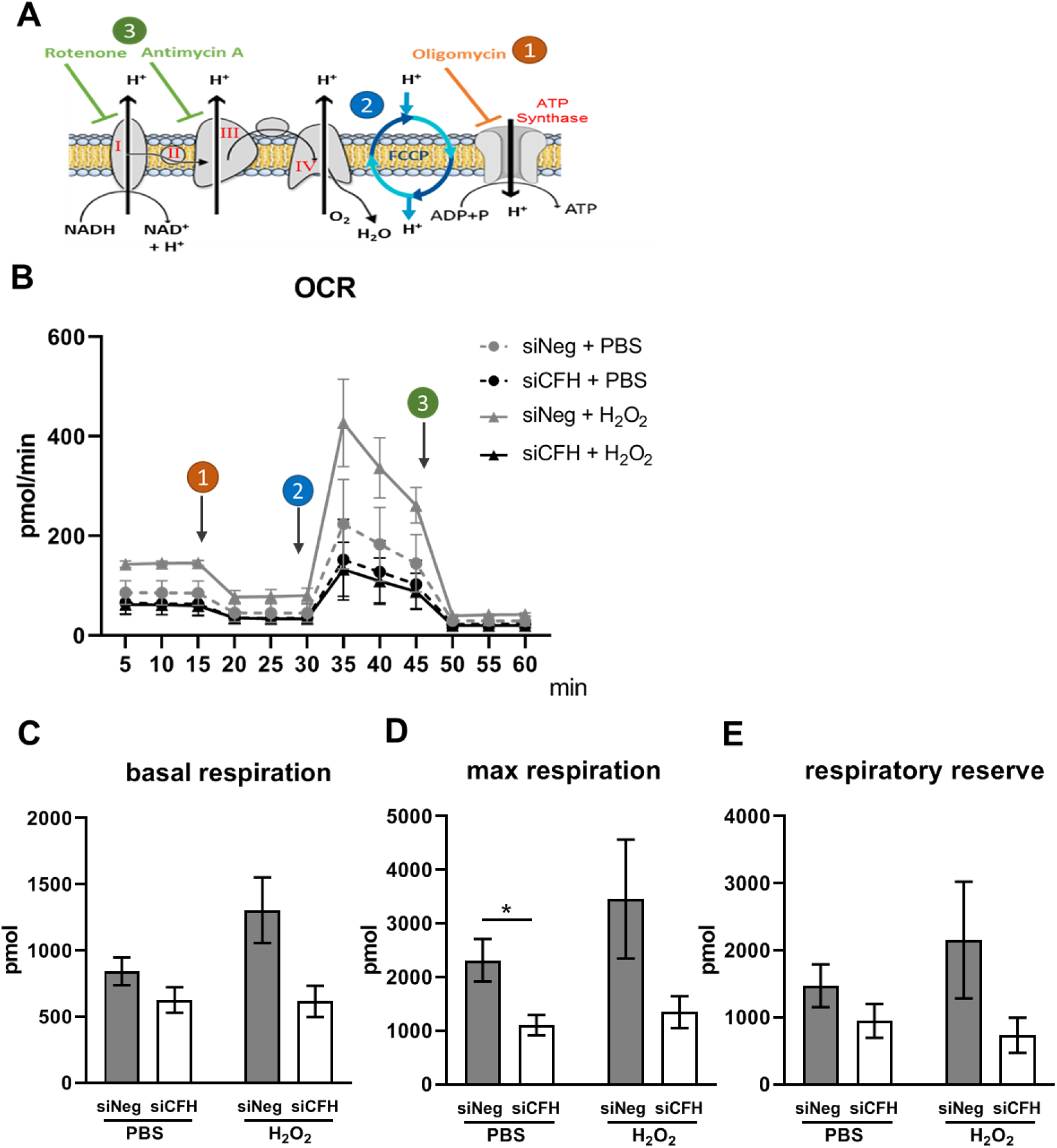
FH loss impairs mitochondrial respiration in RPE cells. **A** Schematic representation of oxidative phosphorylation chain and targeted steps during Seahorse analyses (1,2,3) **B** hTERT-RPE1 cells were seeded, let attach overnight and silenced for 24 hours with negative control (siNeg) or *CFH* specific (si*CFH*) siRNA. 30.000 cells were transferred to seahorse plates overnight and pre-treated for 90 minutes with 200 µM H_2_O_2_ or PBS. Curves show oxygen consumption rate (OCR) measured after 48 hours in hTERT-RPE1. SEM is shown, n=4-8. Arrows indicate injection of oligomycin (1), FCCP (2) and antimycin and rotenone (3) **C-E** Parameters of mitochondrial function are calculated from data shown in B. basal respiration (C), maximal respiration (D) and respiratory reserve (E). Significance was assessed by Student t-test (single effect) and two-way ANOVA (combined effects) as described in the methods section. * p<0.05.

### FH loss alters the expression of mitophagy and mitochondria dynamics genes

Several factors contribute to the energy metabolism regulation and antioxidant capacity of RPE cells, which often both rely on mitochondria function and stability [6, 23]. Impairments in mitochondrial function can be caused by altered oxidative phosphorylation (OxPhos) chain components as shown for Alzheimer’s disease [24], therefore we investigated by qPCR the expression of OxPhos genes NADH dehydrogenase 4 (ND4), Cytochrome c oxidase subunit 4 (COX4) and mitochondrial encoded ATP synthase 6 (ATP6) respectively components of complex 1, complex 4 and ATP synthase [25] (shown in the schematic in Fig 5A). No significant differences were observed in any of the experimental conditions (Supplementary Fig 2A). Transcription factors promoting mitochondrial biogenesis like Peroxisome Proliferator-Activated Receptor Gamma Coactivator 1-Alpha (PGC1a) and Nuclear Factor, Erythroid 2 Like 2 (NRF2) have been shown to positively influence mitochondria metabolism and antioxidant response [26, 27]. To test whether FH loss leads to a dysregulation of those factors, we analyzed gene expression levels of PGC1a and NRF2 (Supplementary Fig. 2B). No differences were detected for NRF2. On the other hand, PGC1a levels were higher in absence of FH. This result would suggest an improvement in mitochondria function, which was not the case in our model. We also measured transcriptional levels of antioxidant enzymes which are induced by PGC1a [28], like Peroxiredoxin 3 (PRDX3), Catalase (CAT), Glutathione Peroxidase 1 (GPX1). We found no differences in PRDX3, a slight upregulation of CAT and a significant increase in GPX1 in the absence of FH (Supplementary Fig. 2C). These data suggest that RPE cells lacking FH are unsuccessfully trying to respond to an oxidative stress situation. Another mechanism of mitochondrial quality control is mitophagy, a mitochondria specific autophagy [29]. We found a significant alteration in the expression levels of genes regulating mitophagy (Fig 6A), like PTEN induced putative kinase 1 (PINK1) and E3 Ubiquitin-Protein Ligase Parkin (PARKIN) and mitochondria dynamics (Fig 6B), like Dynamin-Related Protein 1 (DRP1) and OPA1 Mitochondrial Dynamin Like GTPase (OPA1) when FH was missing (Fig 6B). An alteration in mitophagy levels and mitochondria dynamics would lead to an increase in damaged mitochondria or altered mitochondria turnover.

**Figure 6.**
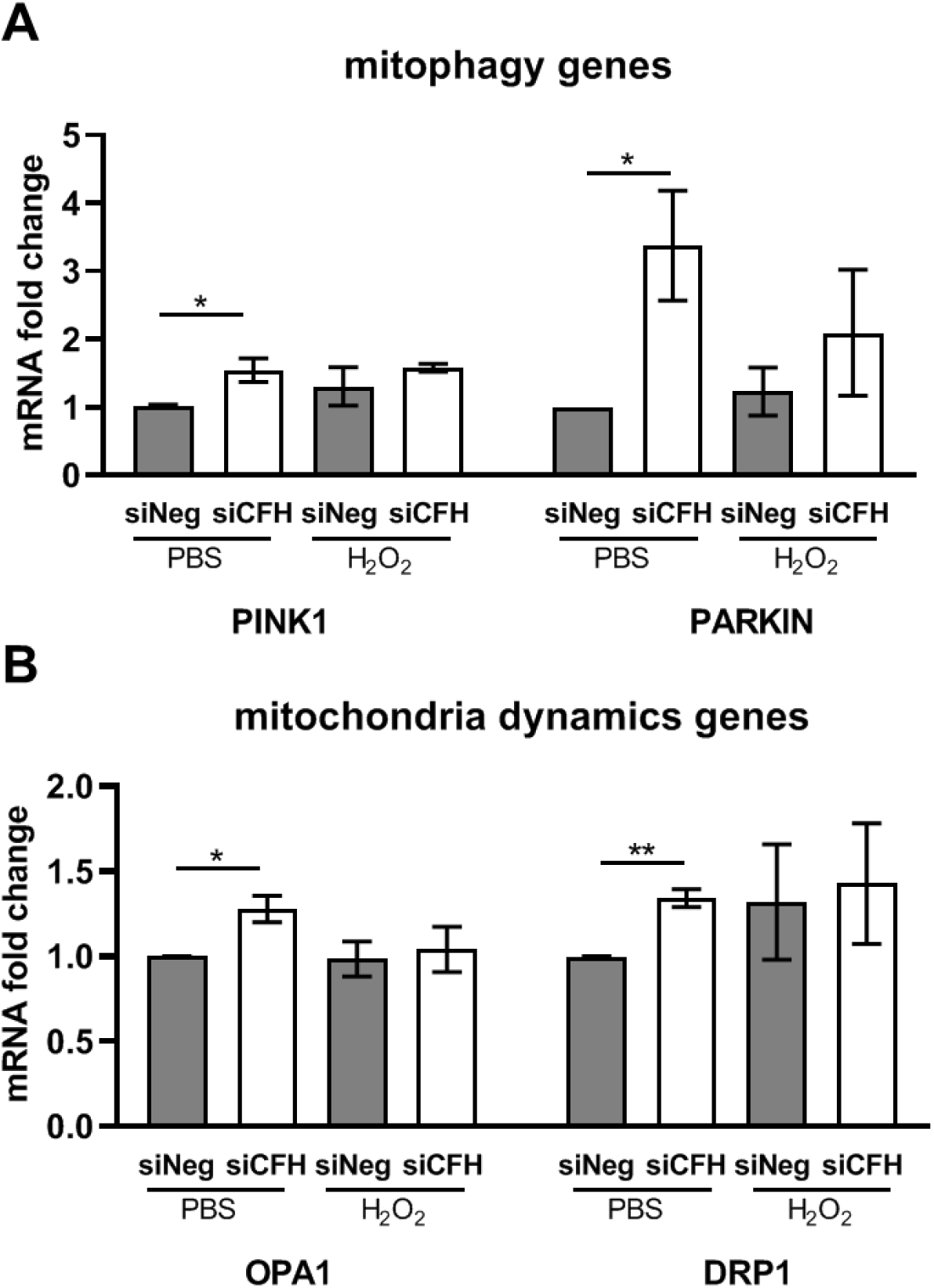
FH modulates expression of mitophagy and mitochondria dynamics genes. hTERT-RPE1 cells were seeded, let attach overnight and silenced for 24 hours with negative control (siNeg) or *CFH* specific (si*CFH*) siRNA. Cells were exposed for 90 minutes to 200 µM H_2_O_2_ or PBS and RNA was collected after 48 hours. **A** gene expression analysis by qRT-PCR of genes involved in mitophagy processes: PTEN Induced Kinase 1 (PINK1) and E3 Ubiquitin-Protein Ligase Parkin (PARKIN) SEM is shown, n=3 **B** gene expression analysis by qRT-PCR of genes involved in mitochondria dynamics: OPA1 Mitochondrial Dynamin Like GTPase (OPA1) and Dynamin-Related Protein 1 (DRP1). SEM is shown, n=3. Data are normalized to housekeeping gene PRPL0 using Δ ΔCt method. Significance was assessed by Student T-test * p<0.05, ** p<0.01.

## Discussion

The retina is a highly organized multi-layered tissue and, in the early stages of AMD, Bruch’s membrane/RPE layer is the first affected. Bruch’s membrane, composed by overlapping extracellular matrix sheets, together with RPE cells separates the retina from the choroid capillary network. RPE cells actively transport nutrients, as glucose, to the retina and eliminate waste material, like oxidized lipids, into the blood stream. Like every other tissue, also the retina undergoes changes with age, including variations in collagen types [30], loss in elastin [31] and increase in metalloproteinases [32]. Bruch’s membrane becomes rigid and thickened, where lipids begin to accumulate underneath the RPE cells [33]. Lipid deposits together with lipofuscin, melanin and complement proteins are the main constituents of drusen, the hallmark lesions associated with AMD pathology [3, 34]. These alterations in Bruch’s membrane structure alter physiological conductivity and therefore impede a correct transport of oxygen or nutrients from the choroidal network to the RPE cells [35-38], resulting in a condition of hypoxia [39] and starvation. In addition, lifestyle habits, like smoking or maintaining a high-fat diet, both risk factors for AMD [7, 40], add oxidative stress to the retina [41, 42], and in particular to the mitochondria of RPE cells and photoreceptors [11]. With age, and more pronounced in AMD patients, mitochondrial damage is augmented [10, 43], thus leading to energy misbalance in the RPE cells. A bioenergetics crisis of RPE cells has been postulated to be at the basis of AMD pathology, especially in relation to the interplay between RPE cells and photoreceptors [6]. Since not the entire elderly population (or the population that smoke) is affected by AMD, other susceptibility factors have to be postulated for RPE cells in AMD patients. Genetic predisposition plays an essential role in AMD pathogenesis and carriers of high-risk variants may not be efficiently responding to ageing processes and oxidative stress. FH risk variants, in particular Y402H, lead to dysregulation of complement activation and hold differential binding properties to oxidized lipids [44, 45]. In this study, we asked to which extent FH dysregulation may alter mechanisms relevant to AMD, like energy metabolism and response to oxidative stress. In the early stages of the disease oxidative stress is limited and can be handled by RPE cells. In addition, Bruch’s membrane is still intact and circulating FH cannot cross choroid/Bruch’s membrane interface due to its size [46]. Therefore, an interplay of genetics and age-related intraretinal changes are likely to drive onset and progression of early AMD. In consequence, we asked whether RPE cells with reduced FH activity are more vulnerable to oxidative insult. The *in vitro* system used in this study allowed us to investigate the effects of endogenous FH specifically on RPE microenvironment without the influence of systemic alterations present in blood. FH dampens activation of the alternative complement pathway. FH acts as cofactor for Complement Factor I (FI), also an inhibitor of complement activation, and displaces Complement Factor B (FB) from C3. Both activities reduce the levels of C3 turnover and complement activation [44]. We show here that RPE cells produce FH, as well as C3. FH downregulation leads to an increase in C3 and C3b levels, prompting complement activation and turnover. In parallel, we detected a slight reduction of FH after H_2_O_2_ pre-treatment, a phenomenon previously observed in H_2_O_2_-induced senescent ARPE19 cells [47]. Besides its role as complement regulator, recent studies implicate FH in lipid metabolism. The AMD-risk-associated FH variant (Y402H) leads in a murine model to a retinal damage similar to AMD and promotes other aspects of the disease, including the accumulation of lipoproteins [48]. Moreover, two FH redox forms have been identified in the circulating blood of AMD patients and those forms hold dual functions [49]. Indeed, the reduced and oxidized forms of FH, as well as FH-Y402 and FH-H402, have different binding affinities to oxidized lipids, which accumulate in drusen [45, 49]. Exogenous FH has been shown to protect ARPE19 cells against H_2_O_2_-induced stress [49] and recently, also against exposure to oxidized lipids 4-HNE (4-hydroxy-2-nonenal) [50]. In this study, we show for the first time a protective role of endogenous *CFH*/FH against oxidative insult. RPE cells, lacking FH, display an accumulation of oxidized lipids in response to H_2_O_2_ pre-treatment. As a consequence, cell viability of RPE cells is affected only in this condition. Lipid oxidation also affects membrane permeability [51] and exogenous FH helps maintain a ZO-1 localization in response to 4-HNE in ARPE19 cells [50]. In our model, we show that reduction of endogenous FH mediates cell membrane damage in RPE cells. Reduction in antioxidant capacity, as well as decrease in viability, could underlie a misbalance in energy metabolism of RPE cells. Several recent evidences show altered bioenergetics being part of AMD pathology [6, 9]. Mitochondria account for the majority of cell energy production, via the tricarboxylic acid (TCA) cycle, OxPhos or lipid breakdown. Nevertheless, mitochondrial metabolism and glycolysis work in a synergetic way, since by-products of glycolysis, like pyruvate, can fuel the TCA. A disruption of either mitochondria or glycolytic function can lead to a failure in the metabolic system. Loss of *CFH* has been associated with mitochondria impairment in retinal development in a *CFH* Knock-out mouse model [52] and patients carrying the *CFH* H402 high-risk variant present increased mitochondrial DNA damage [53]. Whether FH contributes to metabolic homeostasis of RPE cells has never been investigated. We show that FH loss alters energy metabolism and lead to a phenotype similar to the one observed in primary RPE cells derived from AMD patients [9]. After FH reduction, RPE cells show a decline in basal levels of glycolysis and glycolytic capacity, which were both even more affected after pre-treatment with hydrogen peroxide. In the same conditions, GLUT1 expression and glucose uptake were diminished. RPE GLUT1 levels are particularly important for the preservation of the neuroretina. Indeed, in mice with a severe reduction of GLUT1 in RPE cells, glucose transport to the retina was severely hindered and led to photoreceptors cell death [54]. Similar to glycolysis, parameters of mitochondrial respiration were hampered by FH reduction in RPE cells, with the maximal respiration to be the most affected. Interestingly, maximal respiration was increased in control cells after hydrogen peroxide treatment, indicating that RPE cells may respond to a short oxidative insult by increasing their respiration. Of note, this phenomenon was completely abolished in si*CFH* cells. Similar to AMD primary RPE cells, we see in si*CFH* cells upregulation of genes involved in anti-oxidant response, like CAT and GPX1, as well as transcription factors involved in mitochondria stability and biogenesis, like PGC1α. These factors are indicators of a response to oxidative stress and mitochondria damage, but they are not the only ones contributing to define whether a cell will successfully escape from excessive oxidative stress. In fact, we do not observe any improvement in mitochondria function, either in PBS or H_2_O_2_ treated cells, contrarily to AMD primary RPE cells which show a greater resistance toward oxidative stress after 24 hours [9]. This phenomenon may depend on the experimental time. Indeed, another study testing the effect of H_2_O_2_ after 48 hours, a time point used in our model, showed more damage in AMD primary RPE cells after oxidative stress exposure [55]. Cells have developed alternative mechanisms to overcome mitochondrial dysfunction which are activated in case of damage. Mitophagy is a specific type of autophagy directed in eliminating unnecessary, damaged or malfunctioning mitochondria [29]. Mitophagy is classically mediated by the PINK-PARKIN axes [56], and both genes were upregulated in RPE cells when FH levels were reduced. PINK1 accumulates on the membrane of damaged mitochondria and its kinase activity is required for recruitment of PARKIN in order to mediate ubiquitination and organelle removal [56]. PINK1 and PARKIN are also involved in the transport to the lysosomes of mitochondria-derived vesicles (MDV). MDVs contain oxidized proteins which are removed from the mitochondrial matrix and their removal can denote a first effort to rescue mitochondria before engaging in mitophagy [57]. MDV trafficking has been implicated in Parkinson’s and Alzheimer’s disease through association of the vps35 protein, which is mutated in the diseases and involved in MDVs transport [58-60]. In AMD, vesicles accumulation, alterations in autophagy and lysosomes in RPE cells have been described [61, 62], but a role of MDVs in AMD pathology has not yet been considered. PINK1 interacts with proteins involved in mitochondria dynamics including DRP1 and OPA1. Both are fission and fusion genes [63], and were seen upregulated in *CFH* knock-down cells, highlighting the possibility that mitochondrial structures and dynamics are compromised by FH reduction. OPA1 and DRP1 work in concert to maintain mitochondrial stability. Indeed, DRP1 loss-of-function alters OPA1 processing, thus affecting the organization of mitochondrial membranes [64]. Moreover OPA1 is also involved in mitochondrial contraction and inner membrane depolarization, leading to proton leak [65]. Thus, the loss of FH activity likely promotes destabilization of mitochondrial structure and function, followed by perturbation in mitochondrial energy metabolism, structural maintenance of mitochondria and an increase in mitophagy.

In conclusion, this study provides insight into a new mechanism by which FH dysregulation could contribute to processes relevant to AMD. FH reduction renders RPE cells more vulnerable to oxidative stress, with the lipids being particularly affected. RPE cells lacking functional FH show a reduced bioenergetics profile, in regard to both, glycolysis and oxidative phosphorylation. We hypothesized the involvement of mitophagy and mitochondrial dynamics in the process. Taken together, our results suggest a non-canonical role of FH in AMD and highlight its protective role in RPE cells against oxidative stress and metabolic reprogramming, which could help our understanding of the early stages of the disease. Future therapeutic strategies that systemically target the complement system may consider that simple systemic inhibition of complement activity alone may be insufficient to successfully treat AMD.

## Material and Methods

### Cell culture

Human retinal pigment epithelium (RPE) cell line hTERT-RPE1 was obtained from the American Type Culture Collection (ATCC). Cells were maintained in Dulbecco’s modified Eagle’s medium (DMEM; Gibco, Germany) containing 10% fetal calf serum (FCS; Gibco, Germany), penicillin (100 U/mL), streptomycin (100 µg/mL) in a humidified atmosphere containing 5% CO_2_.

### Experimental settings

Cells were seeded in complete growth medium without phenol red in 6- or 24-well plates depending on the experiment and allowed to attach overnight. siRNA mixture with Viromer Blue reagent was prepared according to the manufacturer (Lipocalyx, Germany) using a mix of 3 different double strand hairpin interference RNAs specific for *CFH* and a negative control (Neg), recommended by the provider (IDT technologies, Belgium). In parallel, a positive fluorescent control was used to monitor transfection efficiency (data not shown). Culture medium was substituted with fresh medium and siRNA mixture was added dropwise. After 24 hours, cells were pre-treated for 90 minutes with medium containing 200 µM H_2_O_2_ or PBS as control. Cells were maintained in serum free medium, unless specified otherwise, for the indicated experiment duration. Optimal H_2_O_2_ concentration was assessed in preliminary experiments in hTERT-RPE1 cells, where cell density was monitored using Crystal Violet staining [66] (Supplementary Figure 4). Concentration leading to minimal damage (200 µM) was used for further experiments.

### RNA extraction, cDNA synthesis and quantitative RT-PCR

Cell pellets were collected at the indicated time points and were resolved in 1 ml of TriFAST (PeqLab, Germany), homogenized by inversion and incubated at room temperature for 5 minutes. Then, 200 μl of chloroform was added and the cell pellets vortexed for 15 seconds. Samples were left 10 minutes at room temperature and then centrifuged at 12,000 g for 15 minutes at 4°C. The aqueous phase was transferred into a new tube and mixed with 500 μl of isopropanol for precipitation. After incubation for 10 minutes in ice, samples were centrifuged at 12,000 g for 15 minutes at 4°C. Pellets were rinsed twice with EtOH 75%, dried, resuspended in 20 μl of RNase-free water. RNA purity and concentration were measured using Nanodrop. cDNA was synthesized via reverse-transcription of 2-5 μg of RNA using M-MLV Reverse Transcriptase (200 U, Promega, Wisconsin, USA), random primers (10 ng/μl, Promega, Wisconsin, USA) and dNTPs (0,5 mM) in a total volume of 20 μl. cDNA was used to analyse differences in gene expression by qRT-PCR employing SensiMix SYBR low-Rox KIT (Bioline, Germany) along with gene specific forward and reverse primers (250 nM) listed in Table 1. PCR protocol includes 40 cycles of: 95 °C (15 s), 57 °C (15 s) and 72 °C (25 s). Relative mRNA expression of each gene of interest (GOI) was quantified by using PRPL0 as the housekeeping control gene.

**Table 1.**
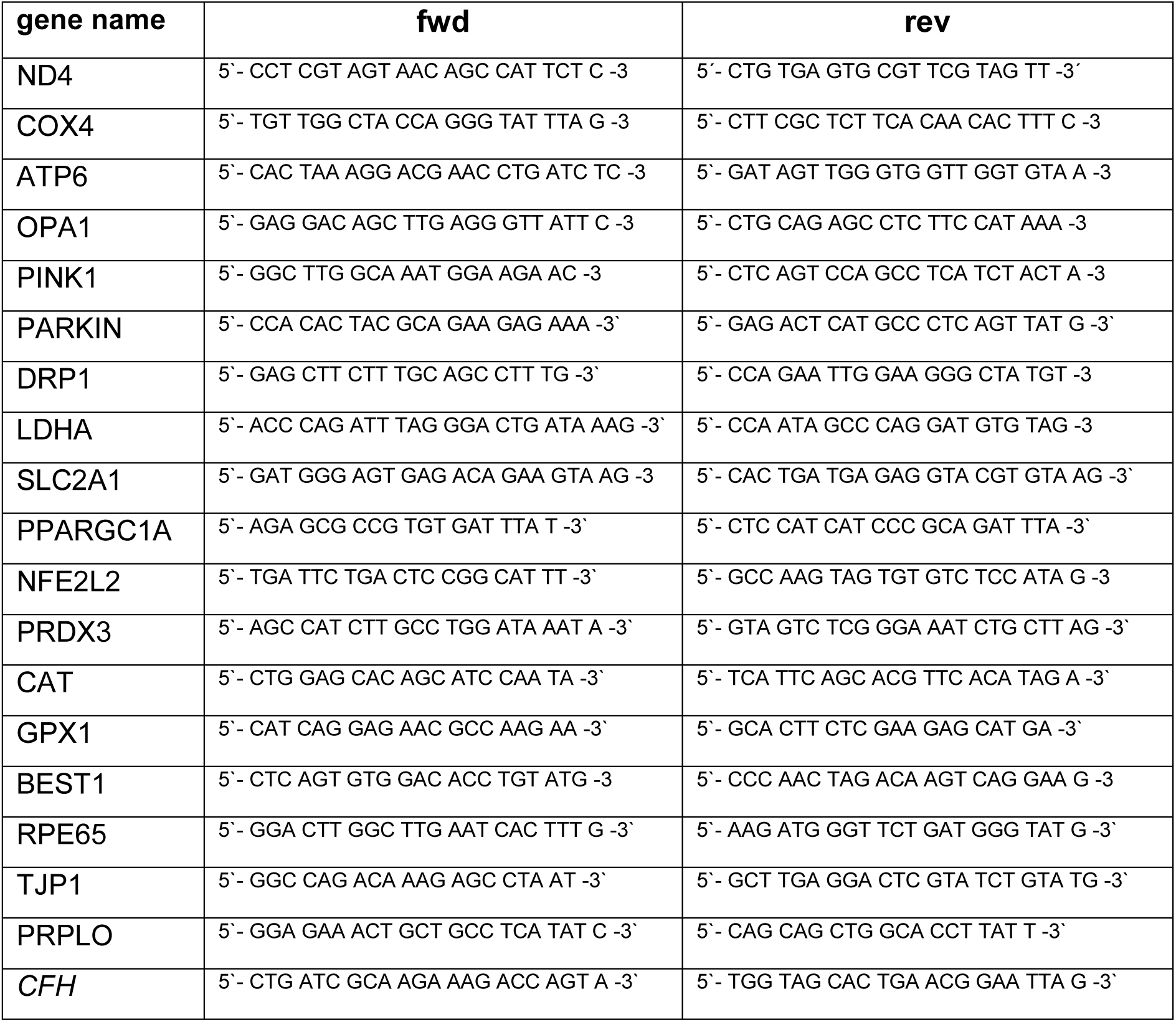
list of qPCR primers

ΔCT=CT(GOI)-CT(housekeeping)

ΔΔCT=ΔCT(sample)-ΔCT(control)

n-fold expression= 2^^-ΔΔCT(GOI)^

### Western Blot

Cell culture supernatants were collected 48 hours after H_2_O_2_ pre-treatment. Following cell debris removal by centrifugation, supernatants were precipitated using ice-cold acetone. Proteins were re-suspended in NuPAGE™ LDS Sample Buffer containing reducing agent (Invitrogen, California, USA), separated on 8-16% or 4-12% SDS-PAGE gels and transferred on PVFD membranes. Membranes were exposed overnight to the primary antibodies (anti-FH, Santa Cruz, Texas, USA; anti-C3, Invitrogen, California, USA) and for 1 hour to HRP-conjugated anti-mouse or anti-rabbit secondary antibody (1:2.000, Cell Signaling, Massachusetts, USA) Immunoreactivity was visualized with Pierce™ ECL Western Blotting Substrate (Thermo Scientific, Massachusetts, USA) and detected with FusionFX instrument (Vilber Lourmat, France).

### C3b ELISA

C3b concentration was evaluated in cell culture supernatants by ELISA assay according to the manufacturer’s instructions (Abcam, UK). Samples were loaded undiluted along with standards and controls in 96 well-plate coated with specific C3b antibody. Absorbance was read at a wavelength of 450 nm immediately after the assay procedure at Spark multimode microplate reader (Tecan, Switzerland). Subtraction readings at 570 nm were taken to correct optical imperfections.

### Cytotoxicity and viability assay

Cytotoxicity and viability were assessed using ApoTox-Glo™ Triplex Assay (Promega, Wisconsin, USA) according to the manufacturer’s instructions. Briefly, two fluorogenic dyes were added to the cell culture media. Viability was assessed by cell-permeable GF-AFC (glycyl-phenylalanyl-aminofluorocoumarin) dye, which is cleaved by live-cell proteases and fluorescence signal is read at 400_Ex_/505_Em_. Cytotoxicity, defined by cell membrane damage, was assessed by cell-impermeable bis-AAF-R110 (bis-alanylalanyl-phenylalanyl-rhodamine 110) dye, which is cleaved by dead-cell proteases released in the cell culture supernatants after membranes damages. Fluorescence is read at 485_Ex_/520_Em_. Spark multimode microplate reader (Tecan, Switzerland) was used for fluorescence measurements. The cleaved products have different excitation/emission readouts; therefore, simultaneous measurements of viability and cytotoxicity were possible. Data are normalized to untreated siNeg controls.

### Lipid peroxidation detection

Lipid peroxidation in live cells was measured via Image-iT® Lipid Peroxidation Kit (Thermo Fischer, Massachusetts, USA), based on BODIPY® 581/591 C11 fluorescent dye. At the indicated time point, the dye was added in cell culture media at the final concentration of 5 µM. Following incubation and washing steps, fluorescence was measured at Spark multimode microplate reader (Tecan, Switzerland). Upon oxidation, the reagent shifts fluorescence emission peak from ∼590 nm to ∼510 nm. Data are shown as ratio of oxidized/reduced lipids.

### Mitochondrial respiration

Mitochondrial function was assessed in live cells using an XFp Extracellular Flux Analyzer (Agilent Technologies, California, USA). After 24 hours silencing (siNeg vs si*CFH*) in 6-well-plates, hTERT-RPE1 cells (3 × 10^4^ cells/well) were seeded in at least duplicates in XFpSeahorse microplates and allowed to adhere overnight. Cells were pre-treated for 90 minutes with medium containing 200 µM H_2_O_2_ or PBS. Following medium change, cells were grown for further 48 hours. Measurements of oxygen consumption rate (OCR) were performed in freshly prepared assay medium, pH 7.4 (Cell Mito Stress Test Assay Medium), according to the manufacturer’s protocol. Mitochondrial function was evaluated after serial injections of 10 μM oligomycin, 10 μM carbonyl cyanide p-trifluoromethoxyphenylhydrazone (FCCP) and 2 µM Antimicyn A / 1 µM Rotenone (all Sigma-Aldrich; Missouri, USA). The data were analyzed using Wave 2.6 Software. All measurements from different experiments were normalized to the mean of the baseline values of siNeg controls.

### Glycolysis

Glycolysis function was assessed in live cells using an XFp Extracellular Flux Analyzer (Agilent Technologies, California, USA). After 24 hours silencing (siNeg vs si*CFH*) in 6-well-plates, hTERT-RPE1 cells (3 × 10^4^ cells/well) were seeded in at least duplicates in XFpSeahorse microplates and allowed to adhere overnight. Cells were pre-treated for 90 minutes with medium containing 200 µM H_2_O_2_ or PBS. Following medium change, cells were grown for further 48 hours. Measurements of extra-cellular acidification rate (ECAR) were performed in freshly prepared assay medium, pH 7.4 (Glycolysis Stress Test Assay Medium), according to the manufacturer’s protocol. Glycolysis was assessed by serially injecting 10 mM Glucose, 10 μM Oligomycin and 50 mM 2-Deoxy-D-Glucose (all Sigma-Aldrich). The data were analyzed using Wave 2.6 Software. All measurements from different experiments were normalized to the mean of the baseline values of siNeg controls.

### Glucose uptake

Glucose uptake was assessed using Glucose Uptake-Glo Assay (Promega, Wisconsin, USA) according to the manufacturer’s instructions. Briefly, at the desired time point cells were washed with PBS and incubated 10 minutes with 1 mM glucose analogue 2-deoxyglucose (2DG), which can be phosphorylated into 2-deoxyglucose-6-phosphate (2DG6P), but not furthered processed by the glycolysis enzymes. After addition of Stop Buffer and neutralization Buffer, 2DG6P Detection reagent containing glucose-6-phosphate dehydrogenase (G6PDH), NADP+, reductase, Glo-luciferase and luciferin substrate was added to allow detection. Luminescence was recorded using Spark multimode microplate reader (Tecan, Switzerland).

### Statistical analysis

All data sets were tested for normal distribution. The combined effects of FH reduction and peroxide pre-treatment on lipid peroxidation, viability; cytotoxicity, glucose uptake and gene expression were assessed with two-way analyses of variance (ANOVA). An unpaired Student T-test was used to compare data from control cells (siNeg) versus si*CFH* cells and H_2_O_2_-treated cells, as well as to compare siNeg and si*CFH* cells after pre-treatment with H_2_O_2_. ELISA data were analyzed using a paired Student t-test. Bioenergetics data were subjected to outliers’ identification via ROUT method. An unpaired Student T-test (or Mann-Whitney test in case the dataset did not pass a normality test) was used to compare data from control cells (siNeg) versus si*CFH* cells and H_2_O_2_-treated cells, as well as to compare siNeg and si*CFH* cells after pre-treatment with H_2_O_2_. Combined effects of FH reduction and peroxide pre-treatment on metabolic parameters were analyzed using two-way ANOVA. Analyses were performed using GraphPad Prism 8 software. Western Blot images were analyzed for signal quantification using Fiji (ImageJ). Significance level was set at p < 0.05.

## Supplementary Figures

**Suppl. Figure 1.**
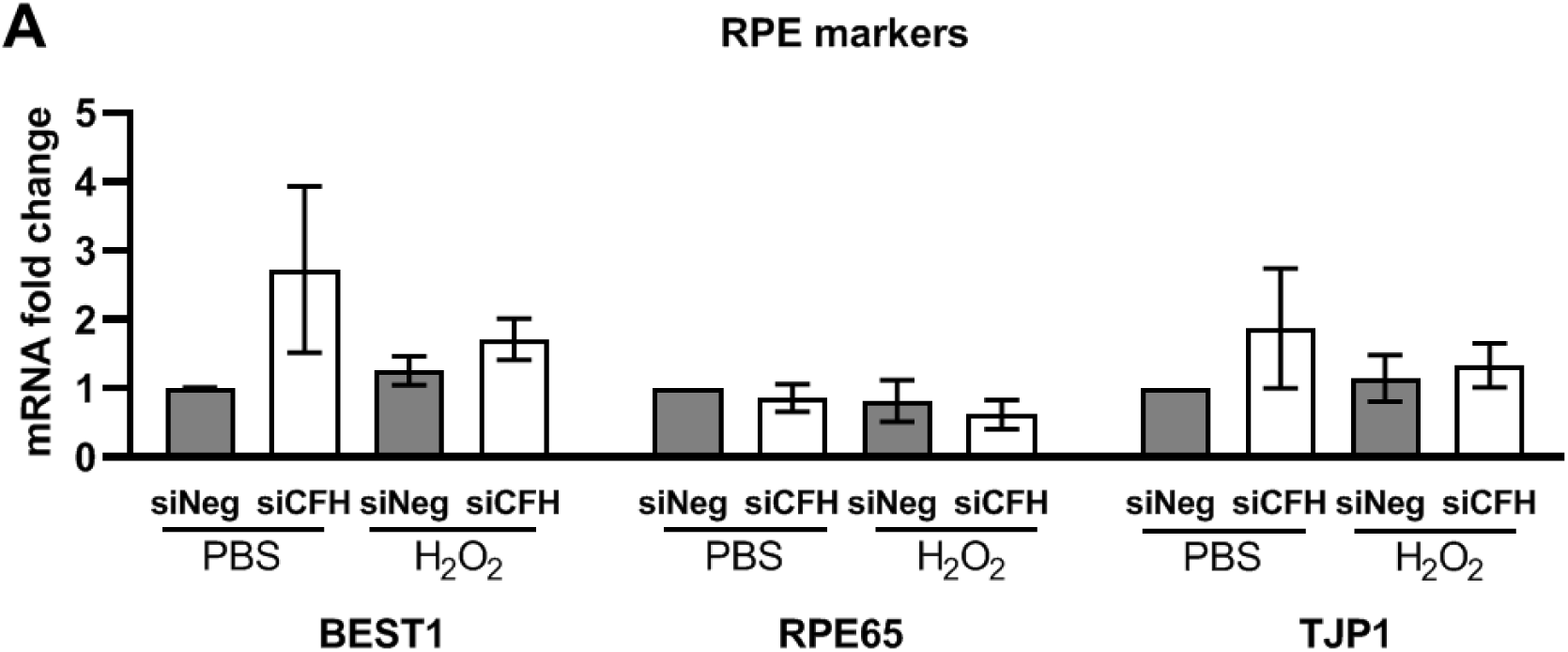
Expression of RPE markers in experimental conditions. Cells were seeded, let attach overnight and silenced for 24 hours with negative control (siNeg) or *CFH* specific (si*CFH*) siRNA. Cells were exposed for 90 minutes to 200 µM H_2_O_2_ or PBS and after 48 hours RNA was collected. **A** Gene expression analysis by qRT-PCR of RPE markers: Bestrophin 1 (BEST1), Retinoid Isomerohydrolase (RPE65), Tight junction protein ZO-1 (TJP1). SEM is shown, n=3. Data are normalized to housekeeping gene PRPL0 using Δ ΔCt method.

**Suppl. Figure 2.**
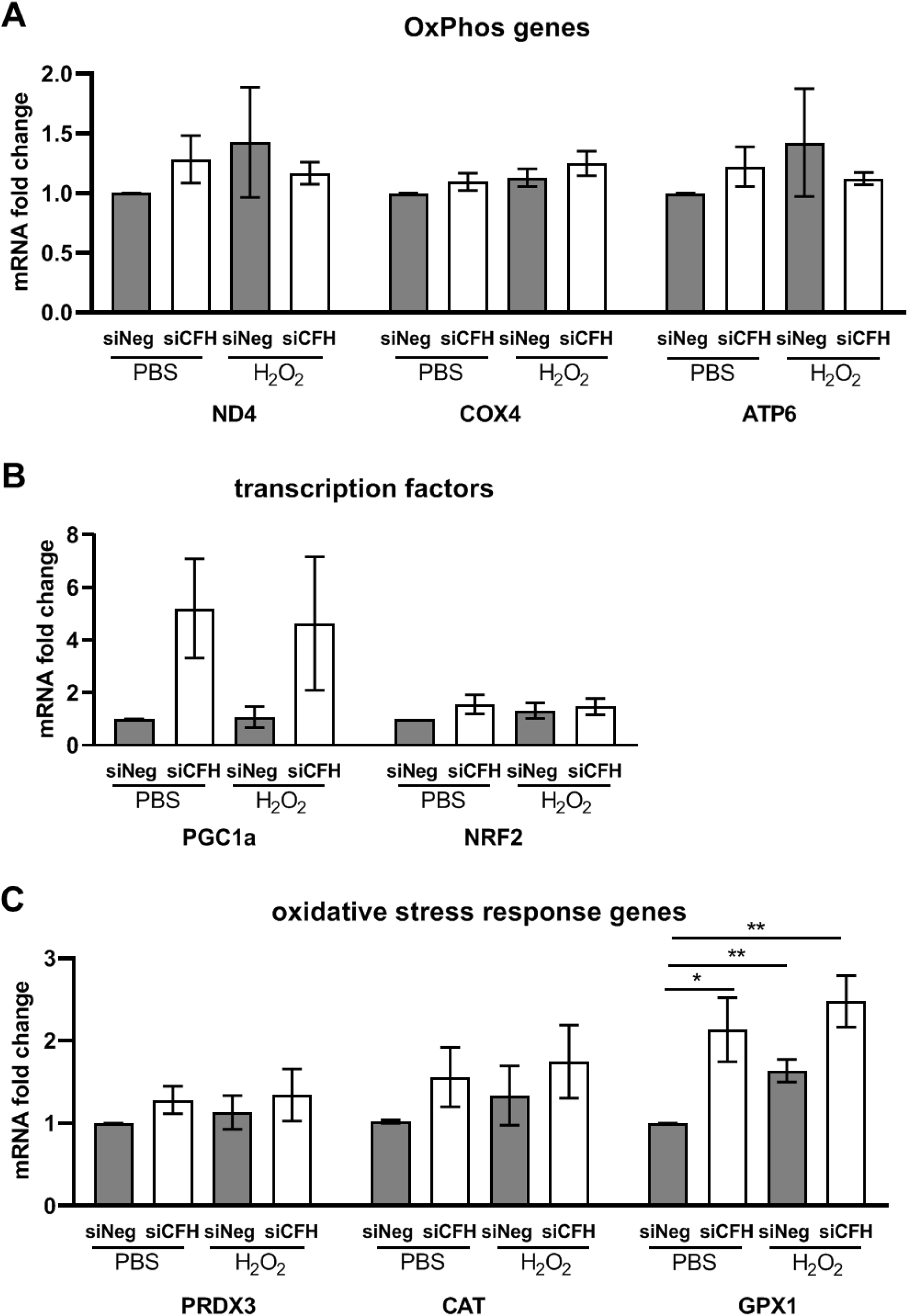
Expression of OxPhos genes, transcription factors and oxidative response genes in experimental conditions. hTERT-RPE1 cells were seeded, let attach overnight and silenced for 24 hours with negative control (siNeg) or *CFH* specific (si*CFH*) siRNA. Cells were exposed for 90 minutes to 200 µM H_2_O_2_ or PBS and after 48 hours RNA was collected. **A** Gene expression analysis by qRT-PCR of OxPhos genes: NADH dehydrogenase 4 (ND4), Cytochrome c oxidase subunit 4 (COX4) and mitochondrially encoded ATP synthase 6 (ATP6). SEM is shown, n=3 **B** Gene expression analysis by qRT-PCR of transcription factors: Peroxisome Proliferator-Activated Receptor Gamma Coactivator 1-Alpha (PGC1a/PPARGC1A) and Nuclear Factor, Erythroid 2 Like 2 (NRF2/NFE2L2). SEM is shown, n=3 **C** gene expression analysis by qRT-PCR of genes involved in oxidative stress response: peroxiredoxin 3 (PRDX3), catalase (CAT), Glutathione Peroxidase 1 (GPX1). SEM is shown, n=3. Data are normalized to housekeeping gene PRPL0 using Δ ΔCt method. Significance was assessed by Student t-test (single effect) and two-way ANOVA (combined effects) as described in the methods section. * p<0.05, ** p<0.01.

**Suppl. Figure 4.**
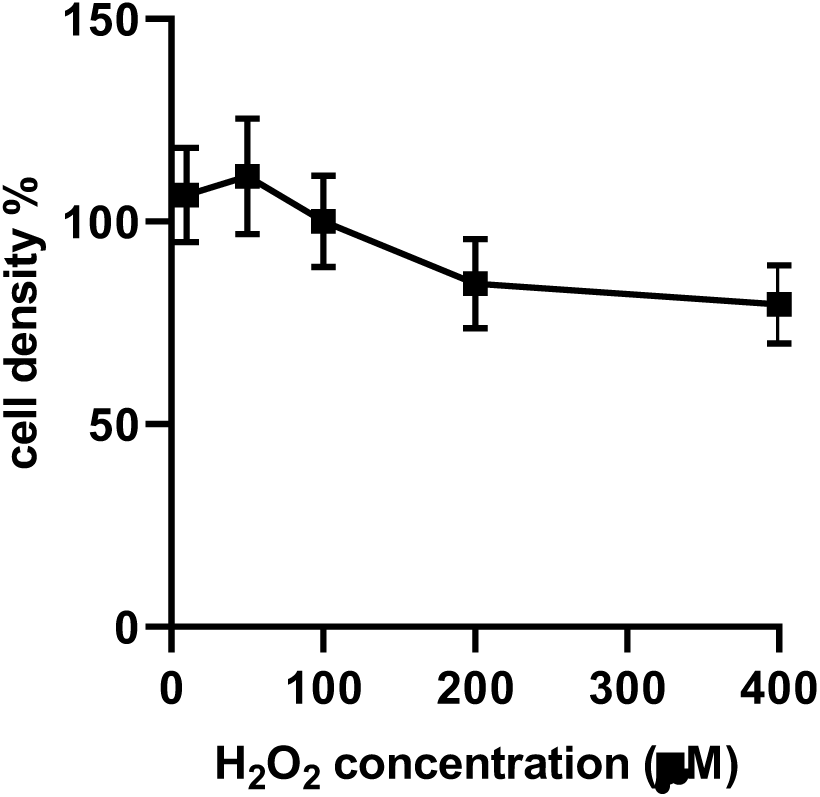
Assessment of optimal concentration for H_2_O_2_ pre-treatment. hTERT-RPE1 cells were seeded, let attach overnight and exposed for 90 minutes to increasing concentrations of H_2_O_2_ and after 48 hours cell density was analyzed via Crystal Violet staining.

**Suppl. Figure 5.**
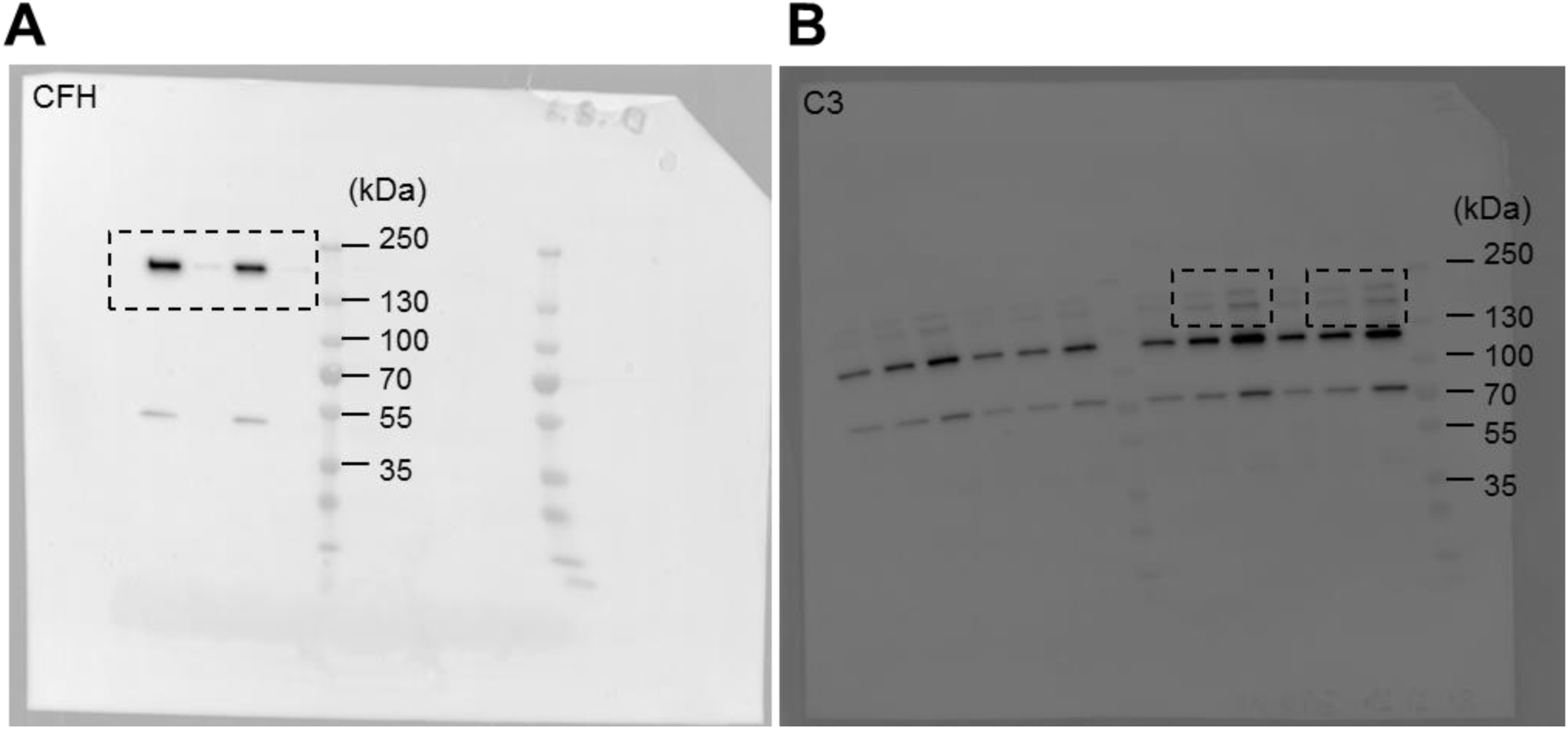
Full images of Western Blots in main Figure 1. A corresponds to Fig 1B. B corresponds to Fig 1D.

## References

1. Wong, W.L., et al., Global prevalence of age-related macular degeneration and disease burden projection for 2020 and 2040: a systematic review and meta-analysis. Lancet Glob Health, 2014. 2(2): p. e106–16.

2. Ferris, F.L., 3rd, et al., Clinical classification of age-related macular degeneration. Ophthalmology, 2013. 120(4): p. 844–51.

3. Rudolf, M., et al., Prevalence and morphology of druse types in the macula and periphery of eyes with age-related maculopathy. Invest Ophthalmol Vis Sci, 2008. 49(3): p. 1200–9.

4. Eamegdool, S.S., et al., Extracellular matrix and oxidative stress regulate human retinal pigment epithelium growth. Free Radic Biol Med, 2019.

5. Strauss, O., The retinal pigment epithelium in visual function. Physiol Rev, 2005. 85(3): p. 845–81.

6. Fisher, C.R. and D.A. Ferrington, Perspective on AMD Pathobiology: A Bioenergetic Crisis in the RPE. Investigative ophthalmology & visual science, 2018. 59(4): p. AMD41–AMD47.

7. van Leeuwen, E.M., et al., A new perspective on lipid research in age-related macular degeneration. Prog Retin Eye Res, 2018. 67: p. 56–86.

8. Chakravarthy, U., et al., Cigarette smoking and age-related macular degeneration in the EUREYE Study. Ophthalmology, 2007. 114(6): p. 1157–63.

9. Ferrington, D.A., et al., Altered bioenergetics and enhanced resistance to oxidative stress in human retinal pigment epithelial cells from donors with age-related macular degeneration. Redox Biol, 2017. 13: p. 255–265.

10. Nordgaard, C.L., et al., Mitochondrial proteomics of the retinal pigment epithelium at progressive stages of age-related macular degeneration. Invest Ophthalmol Vis Sci, 2008. 49(7): p. 2848–55.

11. Brown, E.E., et al., Mitochondrial oxidative stress in the retinal pigment epithelium (RPE) led to metabolic dysfunction in both the RPE and retinal photoreceptors. Redox Biol, 2019. 24: p. 101201.

12. Fritsche, L.G., et al., Age-related macular degeneration: genetics and biology coming together. Annu Rev Genomics Hum Genet, 2014. 15: p. 151–71.

13. Fontaine, M., et al., Truncated forms of human complement factor H. Biochem J, 1989. 258(3): p. 927–30.

14. Edwards, A.O., et al., Complement Factor H Polymorphism and Age-Related Macular Degeneration. Science, 2005. 308(5720): p. 421.

15. Skerka, C., et al., Defective complement control of factor H (Y402H) and FHL-1 in age-related macular degeneration. Mol Immunol, 2007. 44(13): p. 3398–406.

16. Weismann, D., et al., Complement factor H binds malondialdehyde epitopes and protects from oxidative stress. Nature, 2011. 478(7367): p. 76–81.

17. Molins, B., et al., Complement factor H binding of monomeric C-reactive protein downregulates proinflammatory activity and is impaired with at risk polymorphic CFH variants. Sci Rep, 2016. 6: p. 22889.

18. Jun, S., et al., The impact of lipids, lipid oxidation, and inflammation on AMD, and the potential role of miRNAs on lipid metabolism in the RPE. Exp Eye Res, 2018.

19. Datta, S., et al., The impact of oxidative stress and inflammation on RPE degeneration in non-neovascular AMD. Prog Retin Eye Res, 2017. 60: p. 201–218.

20. Nesargikar, P.N., B. Spiller, and R. Chavez, The complement system: history, pathways, cascade and inhibitors. Eur J Microbiol Immunol (Bp), 2012. 2(2): p. 103–11.

21. Kurz, T., et al., ARPE-19 retinal pigment epithelial cells are highly resistant to oxidative stress and exercise strict control over their lysosomal redox-active iron. Autophagy, 2009. 5(4): p. 494–501.

22. Valvona, C.J., et al., The Regulation and Function of Lactate Dehydrogenase A: Therapeutic Potential in Brain Tumor. Brain Pathol, 2016. 26(1): p. 3–17.

23. Barot, M., M.R. Gokulgandhi, and A.K. Mitra, Mitochondrial dysfunction in retinal diseases. Curr Eye Res, 2011. 36(12): p. 1069–77.

24. Manczak, M., et al., Differential expression of oxidative phosphorylation genes in patients with Alzheimer’s disease: implications for early mitochondrial dysfunction and oxidative damage. Neuromolecular Med, 2004. 5(2): p. 147–62.

25. Castellanos, E. and N.J. Lanning, Phosphorylation of OXPHOS Machinery Subunits: Functional Implications in Cell Biology and Disease. Yale J Biol Med, 2019. 92(3): p. 523–531.

26. Kaarniranta, K., et al., PGC-1alpha Protects RPE Cells of the Aging Retina against Oxidative Stress-Induced Degeneration through the Regulation of Senescence and Mitochondrial Quality Control. The Significance for AMD Pathogenesis. Int J Mol Sci, 2018. 19(8).

27. Felszeghy, S., et al., Loss of NRF-2 and PGC-1alpha genes leads to retinal pigment epithelium damage resembling dry age-related macular degeneration. Redox Biol, 2019. 20: p. 1–12.

28. Iacovelli, J., et al., PGC-1alpha Induces Human RPE Oxidative Metabolism and Antioxidant Capacity. Invest Ophthalmol Vis Sci, 2016. 57(3): p. 1038–51.

29. Hyttinen, J.M.T., et al., Mitochondrial quality control in AMD: does mitophagy play a pivotal role? Cell Mol Life Sci, 2018. 75(16): p. 2991–3008.

30. Chen, L., et al., Distribution of the collagen IV isoforms in human Bruch’s membrane. Br J Ophthalmol, 2003. 87(2): p. 212–5.

31. Beattie, J.R., et al., Multiplex analysis of age-related protein and lipid modifications in human Bruch’s membrane. FASEB J, 2010. 24(12): p. 4816–24.

32. Macgregor, A.M., et al., Tissue inhibitor of matrix metalloproteinase-3 levels in the extracellular matrix of lung, kidney, and eye increase with age. J Histochem Cytochem, 2009. 57(3): p. 207–13.

33. Curcio, C.A., et al., Aging, age-related macular degeneration, and the response-to-retention of apolipoprotein B-containing lipoproteins. Prog Retin Eye Res, 2009. 28(6): p. 393–422.

34. Anderson, D.H., et al., The pivotal role of the complement system in aging and age-related macular degeneration: hypothesis re-visited. Prog Retin Eye Res, 2010. 29(2): p. 95–112.

35. Hussain, A.A., L. Rowe, and J. Marshall, Age-related alterations in the diffusional transport of amino acids across the human Bruch’s-choroid complex. J Opt Soc Am A Opt Image Sci Vis, 2002. 19(1): p. 166–72.

36. Hussain, A.A., et al., Macromolecular diffusion characteristics of ageing human Bruch’s membrane: implications for age-related macular degeneration (AMD). Exp Eye Res, 2010. 90(6): p. 703–10.

37. McCarty, W.J., et al., Effects of particulates and lipids on the hydraulic conductivity of Matrigel. J Appl Physiol (1985), 2008. 105(2): p. 621–8.

38. Ramrattan, R.S., et al., Morphometric analysis of Bruch’s membrane, the choriocapillaris, and the choroid in aging. Invest Ophthalmol Vis Sci, 1994. 35(6): p. 2857–64.

39. Arjamaa, O., et al., Regulatory role of HIF-1alpha in the pathogenesis of age-related macular degeneration (AMD). Ageing Res Rev, 2009. 8(4): p. 349–58.

40. Smith, W., et al., Risk factors for age-related macular degeneration: Pooled findings from three continents. Ophthalmology, 2001. 108(4): p. 697–704.

41. Bertram, K.M., et al., Molecular regulation of cigarette smoke induced-oxidative stress in human retinal pigment epithelial cells: implications for age-related macular degeneration. Am J Physiol Cell Physiol, 2009. 297(5): p. C1200–10.

42. Dasari, B., et al., Cholesterol-enriched diet causes age-related macular degeneration-like pathology in rabbit retina. BMC Ophthalmol, 2011. 11: p. 22.

43. Feher, J., et al., Mitochondrial alterations of retinal pigment epithelium in age-related macular degeneration. Neurobiol Aging, 2006. 27(7): p. 983–93.

44. Bajic, G., et al., Complement activation, regulation, and molecular basis for complement-related diseases. EMBO J, 2015. 34(22): p. 2735–57.

45. Shaw, P.X., et al., Complement factor H genotypes impact risk of age-related macular degeneration by interaction with oxidized phospholipids. Proc Natl Acad Sci U S A, 2012. 109(34): p. 13757–62.

46. Clark, S.J., et al., Bruch’s Membrane Compartmentalizes Complement Regulation in the Eye with Implications for Therapeutic Design in Age-Related Macular Degeneration. Frontiers in immunology, 2017. 8: p. 1778–1778.

47. Marazita, M.C., et al., Oxidative stress-induced premature senescence dysregulates VEGF and CFH expression in retinal pigment epithelial cells: Implications for Age-related Macular Degeneration. Redox Biol, 2016. 7: p. 78–87.

48. Landowski, M., et al., Human complement factor H Y402H polymorphism causes an age-related macular degeneration phenotype and lipoprotein dysregulation in mice. Proc Natl Acad Sci U S A, 2019. 116(9): p. 3703–3711.

49. Krilis, M., et al., Dual roles of different redox forms of complement factor H in protecting against age related macular degeneration. Free Radic Biol Med, 2018. 129: p. 237–246.

50. Borras, C., et al., CFH exerts anti-oxidant effects on retinal pigment epithelial cells independently from protecting against membrane attack complex. Sci Rep, 2019. 9(1): p. 13873.

51. Tsubone, T.M., M.S. Baptista, and R. Itri, Understanding membrane remodelling initiated by photosensitized lipid oxidation. Biophys Chem, 2019. 254: p. 106263.

52. Sivapathasuntharam, C., et al., Complement factor H regulates retinal development and its absence may establish a footprint for age related macular degeneration. Sci Rep, 2019. 9(1): p. 1082.

53. Ferrington, D.A., et al., Increased retinal mtDNA damage in the CFH variant associated with age-related macular degeneration. Exp Eye Res, 2016. 145: p. 269–277.

54. Swarup, A., et al., Modulating GLUT1 expression in retinal pigment epithelium decreases glucose levels in the retina: impact on photoreceptors and Muller glial cells. Am J Physiol Cell Physiol, 2019. 316(1): p. C121–C133.

55. Golestaneh, N., et al., Dysfunctional autophagy in RPE, a contributing factor in age-related macular degeneration. Cell Death Dis, 2017. 8(1): p. e2537.

56. Pickrell, A.M. and R.J. Youle, The roles of PINK1, parkin, and mitochondrial fidelity in Parkinson’s disease. Neuron, 2015. 85(2): p. 257–73.

57. Sugiura, A., et al., A new pathway for mitochondrial quality control: mitochondrial-derived vesicles. EMBO J, 2014. 33(19): p. 2142–56.

58. Seaman, M.N., The retromer complex - endosomal protein recycling and beyond. J Cell Sci, 2012. 125(Pt 20): p. 4693–702.

59. Wen, L., et al., VPS35 haploinsufficiency increases Alzheimer’s disease neuropathology. J Cell Biol, 2011. 195(5): p. 765–79.

60. Vilarino-Guell, C., et al., VPS35 mutations in Parkinson disease. Am J Hum Genet, 2011. 89(1): p. 162–7.

61. Keeling, E., et al., Oxidative Stress and Dysfunctional Intracellular Traffic Linked to an Unhealthy Diet Results in Impaired Cargo Transport in the Retinal Pigment Epithelium (RPE). Mol Nutr Food Res, 2019. 63(15): p. e1800951.

62. Sinha, D., et al., Lysosomes: Regulators of autophagy in the retinal pigmented epithelium. Exp Eye Res, 2016. 144: p. 46–53.

63. Yang, Y., et al., Pink1 regulates mitochondrial dynamics through interaction with the fission/fusion machinery. Proc Natl Acad Sci U S A, 2008. 105(19): p. 7070–5.

64. Mopert, K., et al., Loss of Drp1 function alters OPA1 processing and changes mitochondrial membrane organization. Exp Cell Res, 2009. 315(13): p. 2165–80.

65. Lee, H. and Y. Yoon, Transient contraction of mitochondria induces depolarization through the inner membrane dynamin OPA1 protein. J Biol Chem, 2014. 289(17): p. 11862–72.

66. Feoktistova, M., P. Geserick, and M. Leverkus, Crystal Violet Assay for Determining Viability of Cultured Cells. Cold Spring Harb Protoc, 2016. 2016(4): p. pdb prot087379.

